# Dirt floors and domestic animals are associated with soilborne exposure to antimicrobial resistant *E. coli* in rural Bangladeshi households

**DOI:** 10.1101/2025.02.21.639507

**Authors:** Ayse Ercumen, Md. Sakib Hossain, Tahani Tabassum, Ashrin Haque, Amanta Rahman, Md. Hajbiur Rahman, Claire Anderson, Sumaiya Tazin, Suhi Hanif, Gabriella Barratt Heitmann, Md. Rana Miah, Afsana Yeamin, Farjana Jahan, Abul Kasham Shoab, Zahid Hayat Mahmud, Mahbubur Rahman, Jade Benjamin-Chung

## Abstract

Soil can harbor enteropathogens and antimicrobial-resistant organisms in settings with domestic animals. We enrolled 49 households with young children (28 soil floors, 21 concrete floors) in Bangladesh and recorded animal ownership/management. Staff swabbed the floor of children’s sleeping area with a sterile sponge and collected floor dust and a child hand rinse. We used IDEXX QuantiTray/2000 with and without cefotaxime supplementation to enumerate cefotaxime-resistant and generic *E. coli*. There was 8.0 g/m^2^ of dust on soil floors vs. 0.2 g/m^2^ on concrete floors (p-value=0.005). We detected *E. coli* on 100% of soil vs. 86% of concrete floors and cefotaxime-resistant *E. coli* on 89% of soil vs. 43% of concrete floors (p-values<0.05). Cefotaxime-resistant *E. coli* prevalence on floors was 36% in compounds without animals, 79% in compounds with animals and 100% if animals stayed indoors overnight or the floor had animal feces; associations were strongest for chickens. In multivariable models, generic and cefotaxime-resistant *E. coli* counts were 1.5-2 log higher on soil vs. concrete floors, and counts on floors and child hands were 0.17-0.24 log higher for every 10 additional chickens owned (p-values<0.05). Efforts to mitigate infections and antimicrobial resistance in low-income countries should test flooring improvements and hygienic animal management.

**Synopsis:** In rural Bangladeshi households, generic and cefotaxime-resistant *E. coli* were more common on soil floors than concrete floors and among households with higher cohabitation intensity with domestic animals, especially chickens.

## Introduction

Soil in the domestic environment is increasingly recognized as a risk factor for childhood infectious diseases in low-income countries. Young children can ingest soil deliberately via geophagia, in the form of dust, and indirectly via soil-contaminated hands and objects.^1–3^ Soil exposure and geophagia have been linked to adverse child outcomes such as diarrhea,^4,5^ soil-transmitted helminth infections,^6^ environmental enteropathy,^3,7^ and stunting.^7,8^ Enteric pathogens can be deposited on soil surfaces via open defecation and unsafe management of human or animal fecal waste and can have prolonged survival in soil.^9^ In low- and middle-income countries, 42% of household floors are constructed with unimproved materials (e.g., soil, sand, dung, wood, bamboo),^10^ and in Bangladesh, 63% of rural households have soil floors.^11^ Indoor floors made of soil could present a major soilborne exposure pathway to fecal organisms in these settings.

Several studies have investigated the occurrence of fecal indicators in outdoor and indoor domestic soils in low-income countries. In a study in Mozambique, outdoor soil samples collected from house and latrine entrances contained pathogens and molecular markers of human and animal fecal contamination,^12^ and quantitative microbial risk assessment models indicated substantial risk of enteric infections in children from ingestion of fecally contaminated domestic soil.^13^ In our team’s previous study in rural Bangladesh, one gram of outdoor courtyard soil harbored over 120,000 most probable number (MPN) of *Escherichia coli* (*E. coli*),^14^ and higher *E. coli* counts in courtyard soil were associated with higher *E. coli* counts in other domains of the household environment, such as child hands, stored drinking water and stored food.^15^ Fecal indicator bacteria and animal-specific fecal markers have also been commonly detected on indoor household floors in Bangladesh, Peru, Tanzania and Ethiopia.^16–19^

Soil is also a critical reservoir for antimicrobial-resistant organisms and antimicrobial resistance genes because pathogens from human or animal fecal waste deposited on soil surfaces can exchange resistance genes with the high density of native soil microorganisms.^20^ A study in rural Bangladesh found that, among *E. coli* isolates from outdoor courtyard soil, 42% were resistant to at least one antibiotic and 13% were resistant to three or more antibiotics.^21^ In Tanzania, 10% of soil samples from household yards harbored multidrug-resistant *E. coli* ^22^.

Antimicrobial-resistant organisms adapted to soil can then permeate other compartments in the domestic environment.^23^ Studies in both high- and low-income countries have also documented the occurrence of antimicrobial-resistant organisms and antimicrobial resistance genes on indoor floors and in floor dust in public buildings and/or settings with high antimicrobial use (e.g., hospitals, athletic facilities, universities, offices), ^24–28^ as well as in dust inside barns at poultry and pig farms.^29–31^ Studies in the US and Europe have also detected antimicrobial-resistant organisms on residential floors, including bathroom and laundry floors.^32,33^ We are not aware of any studies that have investigated soilborne exposure to antimicrobial-resistant organisms through indoor household floors in low-income countries.

Domestic animals often share living spaces with household members in low-income countries, enabling zoonotic transmission of pathogens and antimicrobial-resistant organisms. Exposure to animals and their feces in the domestic environment is associated with increased risk of child diarrhea^34,35^ and growth deficits.^36,37^ A recent review found that backyard animal husbandry in low-income countries is also associated with exchange of antimicrobial resistance between domestic animals and household members^38^. A sequencing study in Bangladesh demonstrated overlap between human and animal microbiomes and resistomes.^39^ There is growing evidence that domestic soil plays an important role in mediating the transmission of pathogens and antimicrobial-resistant organisms between animals and humans in settings where they commonly cohabitate.^38^ Presence of domestic animals and their fecal waste has been associated with increased abundance of *E. coli* and animal-specific fecal markers in outdoor domestic soil (e.g. courtyard, home entrance) in Bangladesh and Zimbabwe.^15,40,41^ In Ethiopia, fecal indicator levels on indoor floors were higher in households that raised domestic animals and kept them inside during the day.^19^ Also in Ethiopia, *Campylobacter* loads on indoor floors were associated with *Campylobacter* loads in child and chicken feces, indicating potential animal-to-human transmission via floors.^42,43^ Evidence on the links between residential floors, domestic animals and antimicrobial resistance is more limited. In one study of 90 homes in North Carolina, the abundance of the tetracycline resistance gene *tet*(W) in settled indoor dust on the doorsill correlated with the county’s livestock density.^44^ One study in a residential setting in Portugal found overlapping antimicrobial resistance patterns between the inhabitants of a home, their dog and the laundry floor.^33^

Improved flooring materials such as concrete can potentially interrupt soilborne transmission of fecal and/or antimicrobial-resistant microorganisms in low-income countries by providing a floor surface that is easy to clean and does not support microbial survival and growth, and by reducing children’s exposure to soil and dust. A growing body of literature suggests that improved flooring materials are associated with reduced child diarrhea, and protozoan and soil-transmitted helminth infections.^45–49^ Few studies have investigated associations between flooring materials and fecal contamination of floors or other fecal-oral transmission pathways to explore the mechanisms behind these health benefits. In one study in Peru, dirt floors in the house entrance and kitchen areas had significantly higher *E. coli* counts than cement floors in the same areas.^50^ In the same setting, unfinished floors appeared to have higher detection of animal but not human molecular fecal markers than finished floors.^18^ We are not aware of any studies that have investigated associations between floor material and antimicrobial resistance on floors. Further, no studies have explored how floor material and presence of domestic animals jointly affect exposure to fecal and/or antimicrobial-resistant organisms via household floors. Here, we aimed to investigate associations between flooring material, animal ownership and management practices, and detection of *E. coli* and antimicrobial resistant *E. coli* on floors and young children’s hands among households in rural Bangladesh.

## Materials and methods

### Enrollment

We enrolled 49 households with a child <2 years (28 with soil floors, 21 with concrete floors) in rural villages of Sirajganj district in northwestern Bangladesh. The study communities were located in riverine areas that frequently experience flooding. Animal husbandry, specifically cattle rearing, is common in Sirajganj. Anthrax is endemic among ruminants in the area, and outbreaks have been documented among cattle and residents.^51^ To ensure biosafety for field staff when collecting soil samples, households were excluded from enrollment if they reported a case of anthrax among their domestic animals or compound members at any time. The study was approved by the Institutional Review Board of Stanford University (63990) and the Ethical Review Committee of the International Centre for Diarrheoal Disease Research, Bangladesh (icddr,b) (PR-22069).

### Structured questionnaire and spot check observations

In enrolled households, trained field staff administered a structured questionnaire to record animal ownership and management practices, including the number of chickens/ducks, cattle/buffalo, and goats/sheep owned, how frequently these animals were allowed to roam free inside the home or compound, and where the animals were kept at night. Households in rural Bangladesh are typically clustered in extended family compounds; we recorded animal ownership and roaming separately for the enrolled household and for the surrounding compound. Field staff also observed whether any animal feces were present on the floor of the room where the child <2 years slept and observed the materials of the floor, walls and roof.

### Sample collection

Field staff collected floor swabs, dust from floors and (in a systematic subset of 36 households) hand rinse samples from the child <2 years. To collect floor swabs, field staff identified the room where the child slept and used a bleach- and ethanol-sterilized metal stencil to mark a 50 cm x 50 cm square closest to where the child’s head rests when sleeping. They swabbed the area inside the stencil once horizontally and once vertically using a sterile pre-hydrated sponge and placed the sponge in a sterile Whirlpak (Nasco, Modesto, CA). Field staff then marked up to ten additional 50 x 50 cm squares starting from closest to the swabbed area. For each square, they swept the area inside the stencil once horizontally and once vertically using a clean (non-sterile) unused brush and scooped the floor dust into a pre-weighed sterile Whirlpak using a sterile scoop. They recorded how many squares they were able to sweep. To collect hand rinse samples, field staff instructed the caregiver to place the child’s left hand into a sterile Whirlpak pre-filled with 250 mL of sterile deionized water. Once the hand was submerged, they massaged the hand from outside the bag for 15 seconds and then shook the bag with the hand inside for 15 seconds. They repeated the process for the child’s right hand using the same Whirlpak. Ethanol-sterilized gloves were worn for sample collection. Field staff collected 10% field blanks and duplicates for quality control. Floor swab blanks were collected by removing the pre-hydrated sponge from the Whirlpak and placing it back in. Hand rinse blanks were collected by opening the pre-filled Whirlpak and performing the massaging and shaking steps without a child hand in the bag. Samples were transported on ice to the Laboratory of Environmental Health at icddr,b, preserved at 4°C overnight and processed within 24 hours of collection.

### Sample processing

Lab staff recorded the weight of the Whirlpak containing the collected floor dust and subtracted the pre-recorded weight of the bag. Floor swab samples were eluted from the sponge by adding 100 mL of sterile deionized water to the Whirlpak containing the sponge, massaging the sponge from outside the bag for 15 seconds and then swirling the bag for 15 seconds. The liquid was decanted into a new sterile Whirlpak. The process was repeated a total of three times to generate 300 mL of eluate in the Whirlpak.^16^ Lab staff serially diluted the swab eluate 1:10 and 1:100. They generated two 100 mL aliquots with 1:10 dilution (10 mL of eluate + 90 mL of sterile deionized water) and two 100 mL aliquots with 1:100 dilution (10 mL of 1:10 diluted eluate + 90 mL of sterile deionized water). They generated one 100 mL aliquot for hand rinse samples by diluting 50 mL of sample with 50 mL of sterile deionized water.

### Enumeration of generic and cefotaxime-resistant *E. coli*

Lab staff used IDEXX QuantiTray/2000 with Colilert-18 (IDEXX Laboratories, Westbrook, MA) to enumerate the most probable number (MPN) of *E. coli* in floor swab (1:10 and 1:100 dilutions) and child hand rinse samples. For swab samples, they also enumerated the MPN of cefotaxime-resistant *E. coli* by adding 80 μL of filter-sterilized 5 mg/mL cefotaxime solution to the second set of aliquots (1:10 and 1:100 dilutions). The final cefotaxime concentration was 4 μg/mL in the 100 mL sample aliquot. The antibiotic solution was added once Colilert-18 was fully dissolved.^52^ Lab staff performed 10% lab blanks by processing a 100 mL aliquot of sterile deionized water. Trays were incubated at 35°C for 18 hours. Lab staff counted the number of small and large wells that were positive for *E. coli* (yellow and fluorescent under UV) and determined the MPN of *E. coli* using the conversion table provided by IDEXX Laboratories.

Counts for trays below the lower detection limit (1 MPN/100 mL) were imputed as half the lower detection limit (0.5 MPN), and counts for trays above the upper detection limit (2419.6 MPN/100 mL) were imputed as 2420 MPN. For swab samples, if both dilutions were within the detection limits, we calculated an average count weighted by the volume processed (1 mL or 10 mL of eluate). If the 1:100 dilution was below the lower detection limit, we assigned the count from the 1:10 dilution to the sample. If the 1:10 dilution was above the upper detection limit, we assigned the count from the 1:100 dilution to the sample. If either dilution had *E. coli* detected, we considered the sample positive. *E. coli* MPN counts were expressed per unit floor sampling area (0.25 m^2^) for floor swabs and per two hands for child hand rinses. We also calculated log-10 transformed MPN counts for all sample types. For swab samples, we analyzed MPN counts separately for generic *E. coli* and cefotaxime-resistant *E. coli*. We also calculated the percent relative abundance of cefotaxime-resistant *E. coli* for each sample by dividing the MPN counts from the tray processed with cefotaxime supplementation by the MPN counts from the tray processed without cefotaxime supplementation.

### Statistical analysis

We generated binary variables for whether the household had a soil floor, whether the household or the compound owned any chickens/ducks, cattle/buffalo, goats/sheep or any animal, whether these animals ever roamed free inside the home or in the compound, whether they were kept inside the home at night and whether any cattle/buffalo, chicken/duck, goat/sheep or any animal feces were observed on the floor of the child’s sleeping area. We used Mann-Whitney U-tests to compare dust weight per m^2^ between households with soil vs. concrete floors. We used chi-square (χ^2^) tests to compare the prevalence and Mann-Whitney U-tests to compare the log10-transformed MPN counts of generic and cefotaxime-resistant *E. coli* and the relative abundance of cefotaxime-resistant *E. coli* between households in these categories.

To explore dose-response relationships, we categorized households by counts of generic and cefotaxime-resistant *E. coli* and by the percent abundance of cefotaxime-resistant *E. coli* (none detected, and bottom, middle and top tertiles). We compared the number of animals owned across these categories. We also compared *E. coli* prevalence and counts across categories of animal cohabitation intensity (no animal owned by compound, no animal owned by household, animals owned by household but kept outside at night, animals owned by household and kept inside at night) and categories of frequency of animal roaming inside the home or compound (never, sometimes, always). We used the Cuzick nonparametric test for trends across ordered groups for these comparisons.

To assess combined effects of floor type and animal ownership, we used chi-square tests to compare *E. coli* prevalence and Kruskal-Wallis tests to compare log10-tranformed *E. coli* counts between cross-categories of soil vs. concrete floors and households owning at least one vs. no animal. We used multivariable regression with generalized linear models and Gaussian error distribution to estimate associations between floor type and log10-tranformed *E. coli* counts adjusting for the number of animals owned. We conducted analyses separately for each animal type, and for all three animal types combined (i.e., “any animal”).

## Results and discussion

### Household characteristics

We enrolled households between Aug 2-Oct 4, 2023. Of the 49 enrolled households, 28 (57%) had soil floors and 21 (43%) had concrete floors. The main roof and wall material was tin; 48 (98%) households had roofs made of tin and 1 (2%) of cement/concrete, while 42 (86%) households had walls made of tin, 4 (8%) of cement/concrete and 3 (6%) of jute and bamboo. Compounds contained an average of 1.7 (SD=1.4) households and 6.0 (SD=2.1) residents (1.1 children <2 years, 1.5 children between 2-18 years, 3.4 adults>18 years). We did not collect data on socioeconomic indicators in the current study but in a concurrent larger study our team is conducting in the same area, mothers of young children had completed on average 6.2 years of education, the most common occupation for fathers was laborer, and 59% of households had a monthly household income <100 USD (12,000 Taka).

### Floor type and animal management practices

Of 49 households, 71% (35) owned at least one animal (59% owned chickens/ducks, 49% owned cattle/buffalo, 43% owned goats/sheep), and 78% (38) were located in a compound that owned at least one animal (Table S1). Households owned a mean of 16 animals (SD=27), with a mean of 13 chickens/ducks (SD=24), 2 cattle/buffalo (SD=2) and 2 goats/cows (SD=3) (Table S1). Households with soil floors were more likely to own at least one animal and owned more cows but owned a similar number of chickens, goats and overall animals as households with concrete floors (Table S1). Almost all households that owned chickens/ducks reported that these roam free inside the home or in the compound (Table S2). In contrast, approximately half of households that owned cattle/buffalo reported that these roam inside the home or in the compound, while among households that owned goats/sheep, approximately three quarters reported that these roam inside the home and almost all reported that they roam in the compound (Table S2). Animals were reported to roam free inside the home in 75% of households with soil floors vs. 48% of households with concrete floors (Table S1). Among households that owned animals, at least one type of animal was reported to stay inside the home at night in approximately half of households with soil floors and none of households with concrete floors (Table S1). Animal feces were observed in the child’s sleeping area on 14% of soil floors and none of concrete floors (Table S1).

### E. coli counts

We detected generic *E. coli* on 94% (46/49) of floors at a mean log10-transformed MPN of 3.8 (standard deviation [SD]=1.5) per 0.25 m^2^ and on 53% (19/36) of child hands at a mean log-10 transformed MPN of 1.3 (SD=1.0) per two hands. We detected cefotaxime-resistant *E. coli* on 69% (34/49) of floors at a mean log10-transformed MPN of 2.5 (SD=1.3) per 0.25 m^2^. The mean relative abundance of cefotaxime-resistant *E. coli* on floors was 10.5% (SD=22.9).

### Effects of floor type

There was a mean of 8.0 g (SD=13.1) of dust on soil floors vs. 0.2 g (SD=0.3) on concrete floors per m^2^ (p-value=0.005, Table 1). Generic *E. coli* was detected on 100% of soil floors vs. 86% of concrete floors (p-value=0.04), and counts per unit area were 2-log higher on soil floors (log10-mean=4.7) compared to concrete floors (log10-mean=2.7) (p-value<0.0005, Table 1).

**Table 1.** Dust weight on floors, and prevalence and abundance of generic and cefotaxime-resistant *E. coli* on floor swabs and child hands by floor type.

|  | Soil floor | Concrete floor | p-value <sup>a</sup> |
| --- | --- | --- | --- |
| **Floor sweeps** | **N = 28** | **N = 21** |  |
| Dust weight (g/m<sup>2</sup>) | 8.0 (13.1) | 0.2 (0.3) | 0.005 |
| **Floor swabs** | **N = 28** | **N = 21** |  |
| *Generic E. coli* |  |  |  |
| Prevalence, % (n) | 100.0 (28) | 85.7 (18) | 0.04 |
| Log10 MPN, mean (SD) | 4.7 (1.0) | 2.7 (1.2) | <0.0005 |
| *Cefotaxime-resistant E. coli* |  |  |  |
| Prevalence, % (n) | 89.3 (25) | 42.9 (9) | <0.0005 |
| Log10 MPN, mean (SD) | 3.1 (1.1) | 1.6 (0.9) | <0.0005 |
| Percent abundance <sup>b</sup>, mean (SD) | 11.7 (26.0) | 8.7 (17.5) | 0.13 |
| **Child hands** | **N = 19** | **N = 17** |  |
| *Generic E. coli* |  |  |  |
| Prevalence, % (n) | 57.9 (11) | 47.1 (8) | 0.52 |
| Log10 MPN, mean (SD) | 1.6 (1.1) | 1.0 (0.9) | 0.14 |

MPN: Most probable number, SD: Standard deviation.
<sup>a</sup> p-value obtained from chi2 test for prevalence outcomes and Mann-Whitney U-test for dust weight, log10 MPN, and percent abundance outcomes.
<sup>b</sup> Ratio of MPN counts from IDEXX trays with vs. without cefotaxime supplementation for a given sample.

Cefotaxime-resistant *E. coli* was detected on 89% of soil floors and 43% of concrete floors (p-value<0.0005), and counts per unit area were 1.5-log higher on soil floors (log10-mean=3.1) compared to concrete floors (log10-mean=1.6) (p-value<0.0005, Table 1). The relative abundance of cefotaxime-resistant *E. coli* was approximately 10% on both soil and concrete floors. Generic *E. coli* was detected on 58% of child hands in homes with soil floors and 47% in homes with concrete floors, and counts per two hands appeared higher in homes with soil floors (log10-mean=1.6) than in homes with concrete floors (log10-mean=1.0) but the associations with floor type could not be distinguished from chance (Table 1).

### Effects of animal ownership

Generic *E. coli* was detected on 100% of floors if households owned animals vs. 79% of floors if households did not own any animals (p-value=0.005, Table S3). *E. coli* counts per unit area were approximately 1-log higher in households with animals (log10-mean=4.2) vs. without animals (log10-mean=3.1) (p-value=0.03, Table S4). Households in the highest tertile of *E. coli* counts (range: 4.9-5.9 log10 MPN) on floors owned on average 26.9 animals (21.8 chickens/ducks, 2.8 cattle/buffalo, 2.3 goat/sheep) while those in the lowest tertile (range: 1.5-3.3 log10 MPN) owned 12.7 animals (9.1 chickens/ducks, 1.5 cattle/buffalo, 2.1 goat/sheep) and those with no *E. coli* detected on floors owned no animals (trend test p-value=0.004, Table S5, Fig 1). Results were similar for animal ownership by the compound, and the associations were primarily driven by ownership of chickens/ducks (Tables S3-S5, Fig 1).

**Fig 1.**
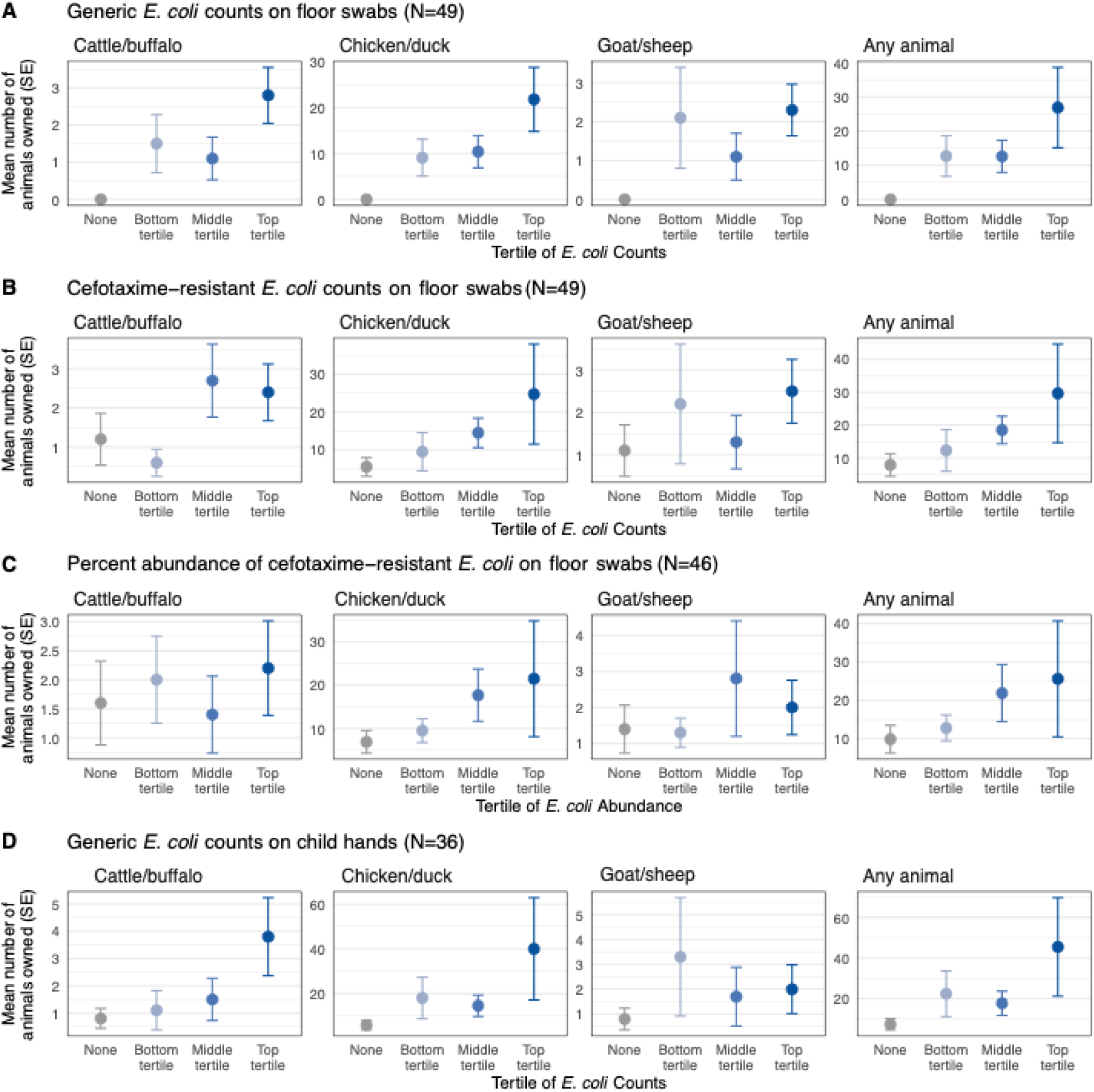
Mean number of animals owned by household by categories of (a) *E. coli* counts on floor swabs, (b) cefotaxime-resistant *E. coli* counts on floor swabs, (c) relative abundance of cefotaxime-resistant *E. coli* on floor swabs, and (d) *E. coli* counts on child hands. *E. coli* counts are categorized as none, and bottom, middle and top tertiles. Circles denote means, and the error bars denote standard errors (SE).

Cefotaxime-resistant *E. coli* was detected on 77% of floors if households owned animals vs. 50% of floors if households did not own animals (p-value=0.06), and on 79% of floors if the compound owned animals vs. 36% of floors if the compound did not own animals (p-value=0.007, Table S3). If the household or compound owned animals, counts of cefotaxime-resistant *E. coli* per unit floor area were approximately 1-log higher than if they did not own animals (p-values<0.05, Table S4). The relative abundance of cefotaxime-resistant *E. coli* on floors was 13% in compounds with animals vs. <1% in compounds without animals (p-value=0.02, Table S4). Households in the highest tertile of cefotaxime-resistant *E. coli* counts (range: 3.5-5.3 log10 MPN) on floors owned on average 29.6 animals (24.7 chickens/ducks, 2.4 cattle/buffalo, 2.5 goat/sheep) while those in the lowest tertile (range: 1.5-2.7 log10 MPN) owned 12.3 animals (9.5 chickens/ducks, 0.6 cattle/buffalo, 2.2 goat/sheep) and those with no cefotaxime-resistant *E. coli* detected on floors owned 7.9 animals (5.5 chickens/ducks, 1.2 cattle/buffalo, 1.1 goats/sheep (trend test p-value=0.01, Table S5, Fig 1). Associations were driven by ownership of chickens/ducks (Tables S3-S5, Fig 1).

Generic *E. coli* was detected on 56-58% of child hands if the household or compound owned animals vs. 42-44% of child hands if the household or compound did not own animals but the differences could not be distinguished from chance (Table S6). *E. coli* counts per two hands were on the order of 1-log and appeared similar in households/compounds with vs. without animals (Table S6). However, children in the highest tertile of *E. coli* counts (range: 2.4-3.8 log10 MPN) on their hands lived in households that owned on average 45.5 animals (40.0 chickens/ducks, 3.8 cattle/buffalo, 2.0 goats/sheep) while those in the lowest tertile (range: 1.0-1.6 log10 MPN) lived in households with 22.4 animals (18.0 chickens/ducks, 1.1 cattle/buffalo, 3.3 goats/sheep) and those with no *E. coli* detected on their hands lived in households with 7.4 animals (5.8 chickens/ducks, 0.8 cattle/buffalo, 0.8 goats/sheep) (trend test p-value=0.06, Table S7, Fig 1).

### Effects of animal management practices

Floors had significantly higher prevalence and counts of generic and cefotaxime-resistant *E. coli* if any animals, specifically chickens, ever roamed free inside the home or in the compound (p-values<0.05, Tables S3-S4). Both the prevalence and counts of generic and cefotaxime-resistant *E. coli* on floors increased progressively as roaming frequency increased from never to sometimes to always (Table S8, Fig 2). Cefotaxime-resistant *E. coli* was detected on 46% of floors (log10-mean=1.9) if animals never roamed in the compound, on 75% of floors (log10-mean=2.6) if they roamed sometimes, and on 100% of floors (log10-mean=3.4) if they always roamed in the compound (trend test p-value=0.02, Table S8, Fig 2). Animal roaming frequency was not associated with *E. coli* prevalence or counts on child hands (Table S9).

**Fig 2.**
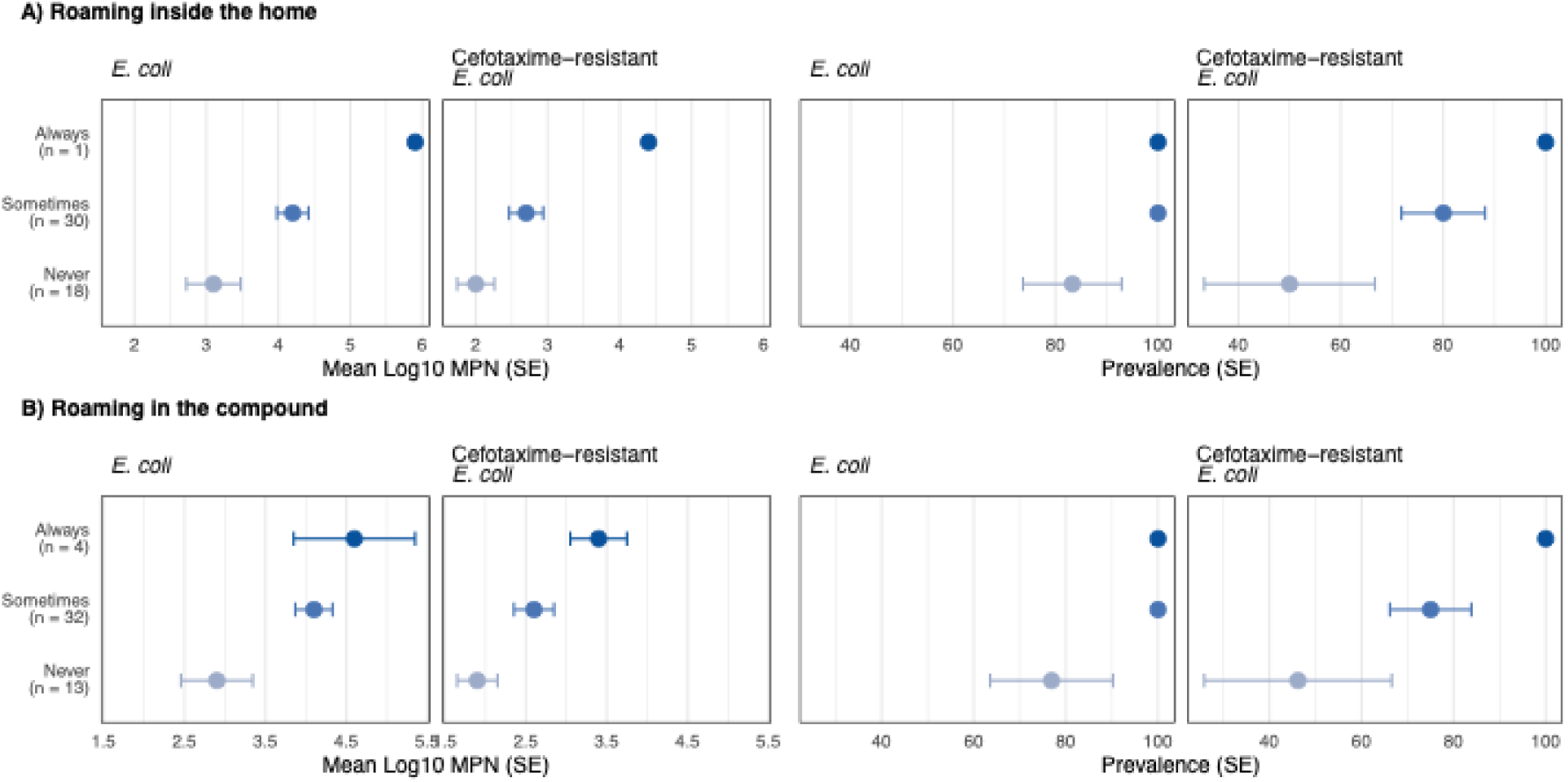
Log-10 transformed most probable number (MPN) and prevalence of generic and cefotaxime-resistant *E. coli* by the free roaming frequency of animals (a) inside the home or (b) in the compound. The compound is a cluster of extended-family households sharing a courtyard. Circles denote means, and the error bars denote standard errors (SE).

Both the prevalence and counts of generic and cefotaxime-resistant *E. coli* on floors also increased progressively as the intensity of animal cohabitation increased (Table S10, Fig 3). Cefotaxime-resistant *E. coli* was detected on 36% of floors (log10-mean=1.8) if the compound did not own any animals, 50% of floors (log10-mean=1.8) if the household did not own any animals, 69% of floors (log10-mean=2.6) if the household owned animals but kept them outside at night, and 100% of floors (log10-mean=3.3) if the household owned animals and kept them indoors at night (trend test p-value=0.01, Table S10, Fig 3). Animal cohabitation intensity was not associated with *E. coli* prevalence or counts on child hands (Table S11).

**Fig 3.**
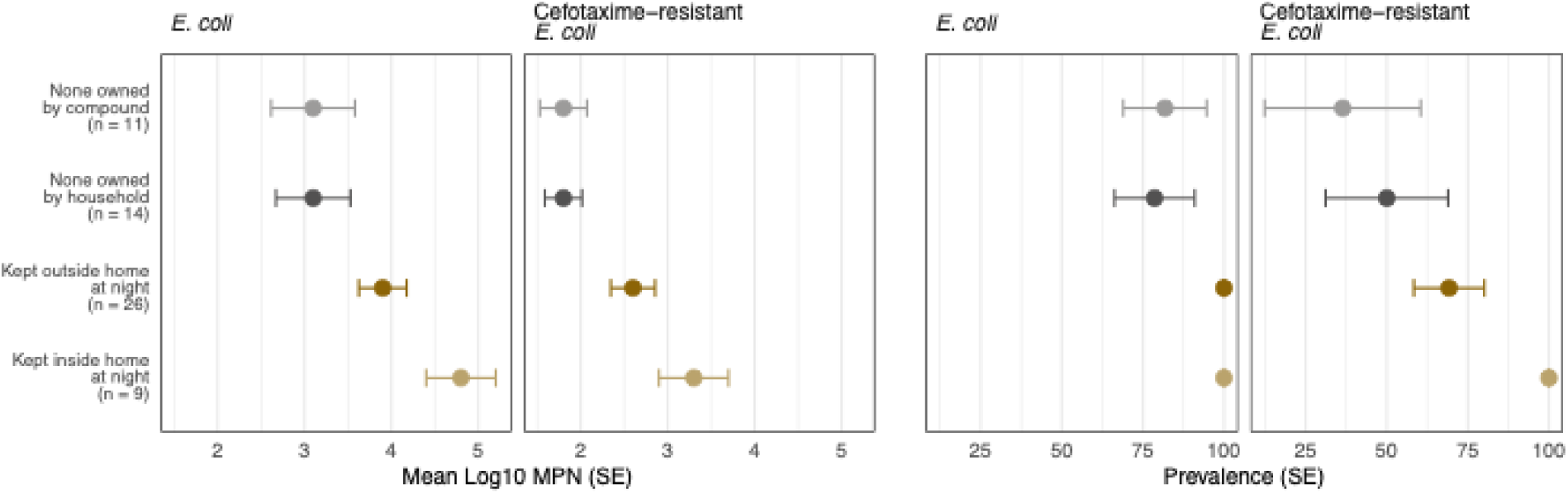
Log-10 transformed most probable number (MPN) and prevalence of generic and cefotaxime-resistant *E. coli* by animal cohabitation intensity. Cohabitation intensity is categorized as no animal owned by compound, no animal owned by household, animals owned by household but kept outdoors at night, and animals owned by household and kept inside the home at night. The compound is a cluster of extended-family households sharing a courtyard. Circles denote means, and the error bars denote standard errors (SE).

Notably, 100% of floors where animal feces were observed harbored generic and cefotaxime-resistant *E. coli* (Table S3). Counts for both were almost 2-log higher on floors with animal feces (log10-mean=5.4 for generic *E. coli*, 4.0 for cefotaxime-resistant *E. coli*) compared to floors without animal feces (log10-mean=3.7 for generic *E. coli*, 2.4 for cefotaxime-resistant *E. coli*) (p-values<0.05, Table S4). Presence of animal feces on the floor was not associated with *E. coli* prevalence or counts on child hands (Table S6).

### Combined effects of floor type and animal ownership

Generic *E. coli* was detected on 63% of concrete floors in households with no animals, vs. 100% of concrete floors in households with animals and 100% of soil floors (p-value=0.001, Table S12, Fig 4). Cefotaxime-resistant *E. coli* was detected on 25% of concrete floors in households with no animals, 54% of concrete floors in households with animals, 83% of soil floors in households without animals, and 91% of soil floors in households with animals (p-value=0.003, Table S12, Fig 4). Both generic and cefotaxime-resistant *E. coli* counts also progressively increased across these cross-categories (p-value=0.001, Table S12, Fig 4). In multivariable regression models controlling for floor type and animal ownership, generic and cefotaxime-resistant *E. coli* counts were 1.5-2 log higher on soil vs. concrete floors, and counts on floor swabs and child hands were 0.13-0.16 log higher for every 10 additional animals owned (p-values<0.05, Table 2). When broken down by animal type, generic *E. coli* counts on floor swabs and child hands and cefotaxime-resistant *E. coli* counts on floor swabs were 0.17-0.24 higher for every 10 additional chickens owned (p-values<0.05); there were no associations with the number of cows and goats owned (Table 2).

**Fig 4.**
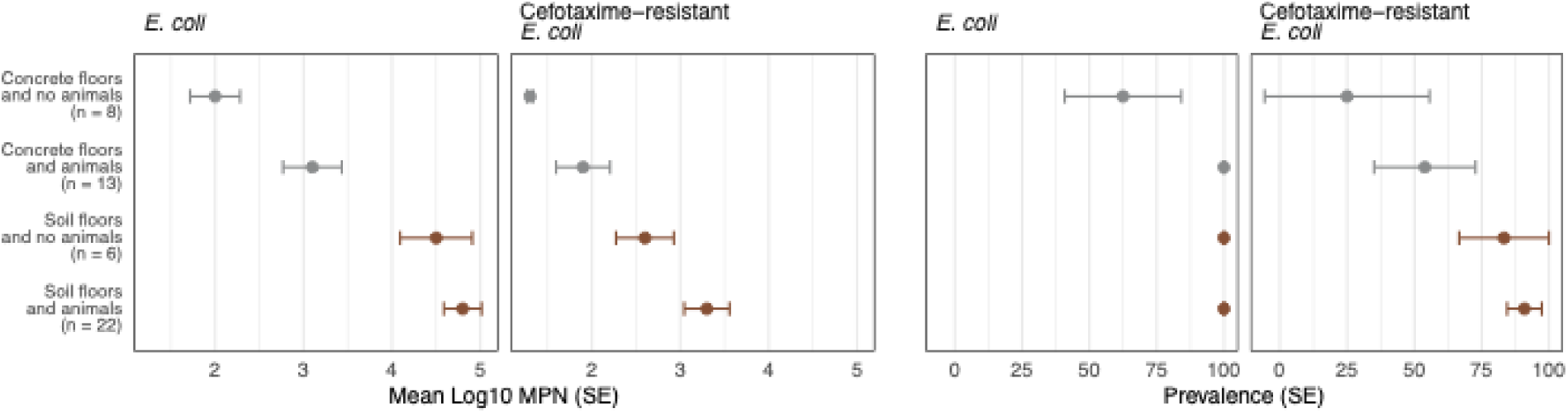
Log-10 transformed most probable number (MPN) and prevalence of generic and cefotaxime-resistant *E. coli* by cross-categories of household floor type and animal ownership. Circles denote means, and the error bars denote standard errors (SE).

**Table 2.** Adjusted associations between household floor type/animal ownership and log10-transformed most probable number (MPN) of *E. coli* and cefotaxime-resistant *E. coli* on floor swabs and *E. coli* on child hands.

|  | Generic *E. coli* on floor swabs |  | Cefotaxime-resistant *E. coli* on floor swabs |  | Generic *E. coli* on child hands |  |
| --- | --- | --- | --- | --- | --- | --- |
| | $\Delta\log_{10}$ (95% CI) | p-value | $\Delta\log_{10}$ (95% CI) | p-value | $\Delta\log_{10}$ (95% CI) | p-value |
| Any animal |  |  |  |  |  |  |
| Soil floor | 2.06 (1.46, 2.67) | <0.0005 | 1.52 (0.97, 2.07) | <0.0005 | 0.57 (-0.94, 1.18) | 0.07 |
| Number of animals<sup>a</sup> | 0.13 (0.01, 0.24) | 0.03 | 0.16 (0.05, 0.26) | 0.003 | 0.14 (0.05, 0.24) | 0.01 |
| By animal type |  |  |  |  |  |  |
| Has soil floor | 2.16 (1.53, 2.79) | <0.0005 | 1.62 (1.04, 2.20) | <0.0005 | 0.49 (-0.13, 1.12) | 0.12 |
| Number of chickens<sup>a</sup> | 0.21 (0.96, 0.36) | 0.01 | 0.24 (0.10, 0.38) | 0.001 | 0.17 (0.02, 0.31) | 0.02 |
| Number of cows | -0.02 (-0.16, 0.12) | 0.79 | -0.01 (-0.15, 0.12) | 0.87 | 0.08 (-0.07, 0.24) | 0.30 |
| Number of goats | -0.08 (-0.19, 0.04) | 0.20 | -0.08 (-0.18, 0.03) | 0.16 | -0.5 (-0.17, 0.06) | 0.34 |

$\Delta\log_{10}$ : Difference in mean log10-transformed MPN counts; CI: Confidence interval.
<sup>a</sup> $\Delta\log_{10}$ reported for every additional 10 animals and every additional 10 chickens

## Discussion

Among 49 households in rural Bangladesh, we found that all soil floors harbored *E. coli* and most (89%) harbored cefotaxime-resistant *E. coli,* while most concrete floors (86%) harbored *E. coli* and 43% harbored cefotaxime-resistant *E. coli.* Soil floors had 40 times more dust per m^2^ than concrete floors. Therefore, the increased detection of *E. coli* and cefotaxime-resistant *E. coli* on soil floors may reflect more dust per unit floor area captured on floor swabs rather than higher microbial abundance per unit mass of dust. However, dustborne exposure to contaminants can be substantial, and therefore, less dust on concrete floors and subsequently fewer fecal/antimicrobial-resistant organisms per unit floor area would translate to reduced child exposure to infectious agents. These findings lend support to a growing body of literature on child health benefits associated with improved floors. Notably, households with soil floors and animals had the highest prevalence and counts of both generic and cefotaxime-resistant *E. coli*, while those with concrete floors and no animals had the lowest. No prior studies have investigated how interactions between floor material and animal ownership and management practices affect contamination of household floors with enteric and/or antimicrobial resistant organisms. Our findings suggest that improved management of domestic animals is critical to achieving the full benefits of concrete floors to prevent the transmission of enteric infections and antimicrobial resistance.

The relative abundance of cefotaxime-resistant *E. coli* on floors was 10 times higher in households that owned animals. The prevalence and counts of both generic and cefotaxime-resistant *E. coli* increased with increasing animal cohabitation intensity (no animals owned by compound, no animals owned by household, animals owned by household but kept outdoors at night, animals owned by household and kept indoors at night) and the frequency of animals roaming free (never, sometimes, always) in the household or compound. Notably, households that had concrete floors were less likely to let their animals roam free inside the home, and no households with concrete floors kept their animals indoors at night or had animal feces observed on floors. A desire to keep concrete floors clean as an aspirational household asset may therefore trigger more hygienic animal management practices as an additional benefit. Our findings are consistent with previous studies that found increased *E. coli* contamination of soil when households owned domestic animals and/or kept them indoors.^43^ Domestic animals are also recognized as reservoirs of antimicrobial resistance due to frequent use of antimicrobials in animal husbandry.^53^ Frequent antibiotic treatment of sick animals, as well as use of antibiotics for healthy animals to promote growth or for prophylaxis is reported among Bangladeshi households that keep domestic animals.^54–56^ A recent review identified soil as an important pathway for transmission of antimicrobial resistance between animals and humans in low-income countries.^38^ A study in Bangladesh found overlapping alleles of antimicrobial resistance genes between humans and domestic animals; chickens had the most overlap with humans.^39^

A study in Kenya identified animal type as an important determinant of the extent of animal contact with, and subsequent risk of contamination on, floors.^57^ Cattle were more likely to be kept in designated sheds both during the day and at night. In contrast, chickens were mostly kept indoors at night, and they roamed freely in and out of the living space during the day, even among households who did not own chickens themselves.^57^ In our analysis, almost all households that owned chickens/ducks, as well as 10-15% of households that did not own chickens/ducks, reported that these roamed freely inside the home or in the compound. In contrast, only half of households reported that cattle roamed freely inside the home or in the compound, approximately 75% reported that goats/sheep roamed inside the home and almost all reported that goats/sheep roamed in the compound. In parallel, we observed stronger relationships between contamination on floors and the presence, number, nighttime location and roaming frequency of chickens/ducks than cattle/buffalo or goats/sheep. In our prior work in rural Bangladesh, chicken feces were most commonly observed (in 87% of compounds vs. cow, goat and sheep feces observed in 19-30% of compounds), and the presence of chickens/chicken feces was more strongly associated with *E. coli* contamination of courtyard soil, as well as child hands, stored water and food than other domestic animals.^15^ In a study in Uganda, ownership of chickens, but not cows, goats or sheep, was associated with higher risk of child diarrhea,^58^ and in Ethiopia, indoor nighttime corralling of chickens, but not other domestic animals, was associated with reduced child growth.^36^ In Nepal, households that owned chickens were more likely to have third-generation cephalosporin-resistant and extended-spectrum beta-lactamase (ESBL)-producing *E. coli* in soil.^59^ Our findings support efforts to improve hygienic management of chickens in low-income settings.^60^

Our study adds to growing evidence that soil floors can be a major source of exposure to antimicrobial-resistant organisms. In our analysis, 89% of soil floors harbored cefotaxime-resistant *E. coli*. Cefotaxime is a third-generation cephalosporin, and cefotaxime resistance is conferred by ESBL production.^61^ Further antibiotic susceptibility testing of 91 cefotaxime-resistant *E. coli* isolates obtained from positive IDEXX trays in our study found that all isolates were multidrug-resistant (resistant to ≥3 antimicrobials), and one isolate was extensively drug-resistant (resistant to ≥12 antimicrobials).^62^ The majority (85%) of isolates contained the *bla_CTX-M_*gene, while 9% of isolates were classified as diarrheagenic and 6% as extraintestinal pathogenic *E. coli* based on the detection of virulence genes.^62^ Additionally, a separate sequencing analysis of soil floor samples in a subset of 10 households from this study detected 45 antimicrobial resistance genes that confer resistance to rifamycin, sulfonamides, tetracyclines, aminoglycosides, among other drug classes.^63^ In a previous study in rural Bangladesh, 42% of *E. coli* isolates from outdoor courtyard soil were antimicrobial-resistant and 13% were multidrug-resistant.^21^ The higher prevalence of antimicrobial-resistant *E. coli* we observed on soil floors may indicate enhanced bacterial survival in the indoor environment. Exposure to sunlight in the outdoor environment can inactivate bacteria; in our previous work in Bangladesh, outdoor soil samples collected from sunlit areas had lower *E. coli* counts than samples collected from shaded areas.^15^ The WHO recommends monitoring ESBL-producing *E. coli* across environmental reservoirs, such as surface waters.^64^ Our findings support also monitoring ESBL-producing *E. coli* in soils.

Our findings indicate that indoor soil floors can present a pathway for soilborne exposure to enteric and antimicrobial-resistant organisms in rural Bangladeshi households, especially when domestic animals share living spaces with humans. Future studies should investigate whether improved floor materials are associated with lower enteric infections and carriage of antimicrobial-resistant organisms among children and household members and evaluate health benefits from safe animal husbandry practices in tandem with flooring improvements.

## Acknowledgement

This study is funded by grants from the National Institute of Child Health and Human Development to Stanford University (R01HD108196).

## Notes

### Competing Interest Statement

The authors have declared no competing interest.

