## Supplemental Information for "Dirt floors and domestic animals are associated with soilborne exposure to antimicrobial resistant *E. coli* in rural Bangladeshi households"


**Table S1.** Animal ownership and management practices 1

**Table S2.** Animal ownership vs. roaming 2

**Table S3.** Prevalence of generic and cefotaxime-resistant *E. coli* on floor swabs by animal ownership and management practices 3

**Table S4.** Log10-transformed most probable number (MPN) of generic and cefotaxime-resistant *E. coli* and relative abundance of cefotaxime-resistant *E. coli* on floor swabs by animal ownership and management practices 4

**Table S5.** Number of animals owned by quartiles of generic and cefotaxime-resistant *E. coli* counts and percent abundance of cefotaxime-resistant *E. coli* on floor swabs 5

**Table S6.** Prevalence and log10-transformed most probable number (MPN) of *E. coli* on child hands by animal ownership and management practices 6

**Table S7.** Number of animals owned by quartiles of *E. coli* counts on child hands 7

**Table S8.** *E. coli* prevalence and abundance on floor swabs by frequency of animal roaming 8

**Table S9.** *E. coli* prevalence and abundance on child hands by frequency of animal roaming 9

**Table S10.** *E. coli* prevalence and abundance on floor swabs by animal cohabitation intensity 10

**Table S11.** *E. coli* prevalence and abundance on child hands by animal cohabitation intensity 11

**Table S12.** *E. coli* prevalence and abundance on floor swabs and child hands by cross-categories of household floor type and animal ownership 12

**Table S1.** Animal ownership and management practices

| | All households (N=49) | Households with soil floors (N=28) | Households with concrete floors (N=21) |
|---|---|---|---|
| **Household owns at least 1, % (n)** | | | |
| Chicken/duck | 59.2 (29) | 71.4 (20) | 42.9 (9) |
| Cattle/buffalo | 49.0 (24) | 57.1 (16) | 38.1 (8) |
| Goat/sheep | 42.9 (21) | 53.6 (15) | 28.6 (6) |
| Any animal | 71.4 (35) | 78.6 (22) | 61.9 (13) |
| **Compound owns at least 1, % (n)** | | | |
| Chicken/duck | 63.3 (31) | 75.0 (21) | 47.6 (10) |
| Cattle/buffalo | 59.2 (29) | 64.3 (18) | 52.4 (11) |
| Goat/sheep | 46.9 (23) | 57.1 (16) | 33.3 (7) |
| Any animal | 77.6 (38) | 82.1 (23) | 71.4 (15) |
| **Number of animals in household, mean (SD)** | | | |
| Chicken/duck | 12.8 (24.0) | 11.5 (12.6) | 14.5 (34.2) |
| Cattle/buffalo | 1.7 (2.4) | 2.3 (2.8) | 0.9 (1.5) |
| Goat/sheep | 1.7 (3.1) | 1.9 (2.3) | 1.5 (3.9) |
| Any animal | 16.2 (26.7) | 15.7 (15.1) | 16.9 (37.4) |
| **Number of animals in compound, mean (SD)** | | | |
| Chicken/duck | 14.1 (24.1) | 13.9 (12.9) | 14.4 (34.2) |
| Cattle/buffalo | 2.2 (2.9) | 2.8 (3.3) | 1.5 (2.1) |
| Goat/sheep | 1.9 (3.1) | 2.0 (2.2) | 1.8 (4.0) |
| Any animal | 18.2 (26.5) | 18.7 (15.4) | 17.6 (36.9) |
| **Animals ever roam free in household:** | | | |
| Chicken/duck | 61.2 (30) | 75.0 (21) | 42.9 (9) |
| Cattle/buffalo | 26.5 (13) | 35.7 (10) | 14.3 (21) |
| Goat/sheep | 34.7 (17) | 50.0 (14) | 14.3 (3) |
| Any animal | 63.3 (31) | 75.0 (21) | 47.6 (10) |
| **Animals ever roam free in compound:** | | | |
| Chicken/duck | 65.3 (32) | 75.0 (21) | 52.4 (11) |
| Cattle/buffalo | 28.6 (14) | 32.1 (9) | 23.8 (5) |
| Goat/sheep | 42.9 (21) | 57.1 (16) | 23.8 (5) |
| Any animal | 73.5 (36) | 82.1 (23) | 61.9 (13) |
| **Animals kept inside home at night [a]:** | | | |
| Chicken/duck | 6.9 (2) | 10.0 (2) | 0.0 (0) |
| Cattle/buffalo | 12.5 (3) | 18.8 (3) | 0.0 (0) |
| Goat/sheep | 38.1 (8) | 53.3 (8) | 0.0 (0) |
| Any animal | 25.7 (9) | 40.9 (9) | 0.0 (0) |
| **Animal feces observed on floor [a]:** | | | |
| Chicken/duck | 4.1 (2) | 7.1 (2) | 0.0 (0) |
| Cattle/buffalo | 4.1 (2) | 7.1 (2) | 0.0 (0) |
| Goat/sheep | 2.0 (1) | 3.6 (1) | 0.0 (0) |
| Any animal | 8.2 (4) | 14.3 (4) | 0.0 (0) |

**Table S2.** Animal ownership vs. roaming

| | Yes | Roams in household | Roams in compound | No | Roams in household | Roams in compound |
|---|---|---|---|---|---|---|
| **Household owns:** | **N** | **% (n)** | **% (n)** | **N** | **% (n)** | **% (n)** |
| Chickens | 29 | 96.6 (28) | 100.0 (29) | 20 | 10.0 (2) | 15.0 (3) |
| Cows | 24 | 45.8 (11) | 50.0 (12) | 25 | 8.0 (2) | 8.0 (2) |
| Goat | 21 | 76.2 (16) | 95.2 (20) | 28 | 3.6 (1) | 3.6 (1) |
| Any animal | 35 | 82.9 (29) | 94.3 (33) | 14 | 14.3 (2) | 21.4 (3) |
| **Compound owns:** | **N** | **% (n)** | **% (n)** | **N** | **% (n)** | **% (n)** |
| Chickens | 31 | 93.6 (29) | 100.0 (31) | 18 | 5.6 (1) | 5.6 (1) |
| Cows | 29 | 44.8 (13) | 48.3 (14) | 20 | 0.0 (0) | 0.0 (0) |
| Goat | 23 | 73.9 (17) | 91.3 (21) | 26 | 0.0 (0) | 0.0 (0) |
| Any animal | 38 | 79.0 (30) | 92.1 (35) | 11 | 9.1 (1) | 9.1 (1) |

**Table S3.** Prevalence of generic and cefotaxime-resistant *E. coli* on floor swabs by animal ownership and management practices

| | | | Generic *E. coli* | | | Cefotaxime-resistant *E. coli* | | |
|---|---|---|---|---|---|---|---|---|
| | N | | Prevalence % (n) | | Chi2 | Prevalence % (n) | | Chi2 |
| | Yes | No | Yes | No | p-value | Yes | No | p-value |
| **Household owns at least 1:** | | | | | | | | |
| Chicken/duck | 29 | 20 | 100.0 (29) | 85.0 (17) | 0.03 | 79.3 (23) | 55.0 (22) | 0.07 |
| Cattle/buffalo | 24 | 25 | 100.0 (24) | 88.0 (22) | 0.08 | 79.2 (19) | 60.0 (15) | 0.15 |
| Goat/sheep | 21 | 28 | 100.0 (21) | 89.3 (25) | 0.12 | 76.2 (16) | 64.3 (18) | 0.37 |
| Any animal | 35 | 14 | 100.0 (35) | 78.6 (11) | 0.005 | 77.1 (27) | 50.0 (14) | 0.06 |
| **Compound owns at least 1:** | | | | | | | | |
| Chicken/duck | 31 | 18 | 100.0 (31) | 83.3 (15) | 0.02 | 83.9 (26) | 44.4 (8) | 0.004 |
| Cattle/buffalo | 29 | 20 | 100.0 (29) | 85.0 (17) | 0.03 | 79.3 (23) | 55.0 (11) | 0.07 |
| Goat/sheep | 23 | 26 | 95.7 (22) | 92.3 (24) | 0.63 | 73.9 (17) | 65.4 (17) | 0.52 |
| Any animal | 38 | 11 | 97.4 (37) | 81.8 (9) | 0.06 | 79.0 (30) | 36.4 (4) | 0.007 |
| **Animals ever roam free in household:** | | | | | | | | |
| Chicken/duck | 30 | 19 | 100.0 (30) | 84.2 (16) | 0.03 | 80.0 (24) | 52.6 (10) | 0.04 |
| Cattle/buffalo | 13 | 36 | 100.0 (13) | 91.7 (33) | 0.28 | 84.6 (2) | 63.9 (23) | 0.17 |
| Goat/sheep | 17 | 32 | 100.0 (17) | 90.6 (29) | 0.19 | 82.4 (14) | 62.5 (20) | 0.15 |
| Any animal | 31 | 18 | 100.0 (31) | 83.3 (15) | 0.02 | 80.7 (25) | 50.0 (9) | 0.03 |
| **Animals ever roam free in compound:** | | | | | | | | |
| Chicken/duck | 32 | 17 | 100.0 (32) | 82.4 (14) | 0.01 | 81.3 (26) | 47.1 (8) | 0.01 |
| Cattle/buffalo | 14 | 35 | 100.0 (14) | 91.4 (32) | 0.26 | 71.4 (25) | 64.3 (9) | 0.62 |
| Goat/sheep | 21 | 28 | 100.0 (21) | 89.3 (25) | 0.12 | 76.2 (16) | 64.3 (18) | 0.37 |
| Any animal | 36 | 13 | 100.0 (36) | 76.9 (10) | 0.003 | 77.8 (28) | 46.2 (6) | 0.03 |
| **Animals kept inside home at night [a]:** | | | | | | | | |
| Chicken/duck | 2 | 27 | 100.0 (2) | 100.0 (2) | -- | 100.0 (2) | 77.8 (21) | 0.62 |
| Cattle/buffalo | 3 | 21 | 100.0 (3) | 100.0 (21) | -- | 100.0 (3) | 76.2 (16) | 0.48 |
| Goat/sheep | 8 | 13 | 100.0 (8) | 100.0 (13) | -- | 100.0 (8) | 61.5 (8) | 0.06 |
| Any animal | 9 | 26 | 100.0 (9) | 100.0 (26) | -- | 100.0 (9) | 69.2 (18) | 0.07 |
| **Animal feces observed on floor [a]:** | | | | | | | | |
| Chicken/duck | 2 | 47 | 100.0 (2) | 93.6 (44) | 0.88 | 100.0 (2) | 68.1 (32) | 0.48 |
| Cattle/buffalo | 2 | 47 | 100.0 (2) | 93.6 (44) | 0.88 | 100.0 (2) | 68.1 (32) | 0.48 |
| Goat/sheep | 1 | 48 | 100.0 (1) | 93.8 (45) | 0.94 | 100.0 (1) | 68.8 (33) | 0.69 |
| Any animal | 4 | 45 | 100.0 (4) | 93.3 (42) | 0.77 | 100.0 (4) | 66.7 (30) | 0.22 |

[a] Fisher's exact test performed because of small cells.

**Table S4.** Log10-transformed most probable number (MPN) of generic and cefotaxime-resistant *E. coli* and relative abundance of cefotaxime-resistant *E. coli* on floor swabs by animal ownership and management practices

| | | | **Generic *E. coli*** | | | **Cefotaxime-resistant *E. coli*** | | | **% abundance** | | |
|---|---|---|---|---|---|---|---|---|---|---|---|
| | **N** | | **Log10 MPN Mean (SD)** | | **MW** | **Log10 MPN Mean (SD)** | | **MW** | **Mean (SD)** | | **MW** |
| | **Yes** | **No** | **Yes** | **No** | **p-value** | **Yes** | **No** | **p-value** | **Yes** | **No** | **p-value** |
| **Household owns at least 1:** | | | | | | | | | | | |
| Chicken/duck | 29 | 20 | 4.3 (1.3) | 3.2 (1.6) | 0.02 | 2.8 (1.3) | 2.0 (1.1) | 0.03 | 13.3 (26.5) | 5.6 (14.4) | 0.26 |
| Cattle/buffalo | 24 | 25 | 4.2 (1.4) | 3.5 (1.5) | 0.07 | 2.8 (1.3) | 2.2 (1.2) | 0.06 | 16.9 (30.1) | 3.5 (1.2) | 0.30 |
| Goat/sheep | 21 | 28 | 4.2 (1.4) | 3.6 (2.5) | 0.14 | 2.8 (1.4) | 2.2 (1.1) | 0.14 | 14.9 (30.4) | 6.8 (13.5) | 0.64 |
| Any animal | 35 | 14 | 4.2 (1.4) | 3.1 (1.6) | 0.03 | 2.8 (1.3) | 1.8 (0.8) | 0.02 | 13.5 (25.6) | 1.1 (1.6) | 0.09 |
| **Compound owns at least 1:** | | | | | | | | | | | |
| Chicken/duck | 31 | 18 | 4.3 (1.2) | 3.1 (1.6) | 0.01 | 2.8 (1.3) | 2.0 (1.1) | 0.03 | 12.7 (25.7) | 6.0 (15.4) | 0.10 |
| Cattle/buffalo | 29 | 20 | 4.2 (1.3) | 3.4 (1.7) | 0.11 | 2.7 (1.3) | 2.3 (1.3) | 0.19 | 14.2 (28.0) | 4.2 (6.0) | 0.54 |
| Goat/sheep | 23 | 26 | 4.1 (1.5) | 3.6 (1.5) | 0.25 | 2.8 (1.4) | 2.2 (1.1) | 0.15 | 15.0 (30.0) | 6.4 (13.6) | 0.43 |
| Any animal | 38 | 11 | 4.1 (1.4) | 3.1 (1.6) | 0.06 | 2.7 (1.3) | 1.8 (0.9) | 0.03 | 12.9 (25.0) | 0.8 (1.6) | 0.02 |
| **Animals ever roam free in household:** | | | | | | | | | | | |
| Chicken/duck | 30 | 19 | 4.3 (1.2) | 3.1 (1.6) | 0.01 | 2.8 (1.3) | 2.0 (1.1) | 0.03 | 12.8 (26.2) | 6.2 (14.8) | 0.42 |
| Cattle/buffalo | 13 | 36 | 4.3 (1.2) | 3.7 (1.6) | 0.25 | 2.8 (1.3) | 2.4 (1.3) | 0.31 | 20.5 (36.7) | 6.6 (13.3) | 0.38 |
| Goat/sheep | 17 | 32 | 4.4 (1.3) | 3.5 (1.5) | 0.04 | 3.1 (1.4) | 2.2 (1.1) | 0.04 | 18.1 (33.2) | 6.0 (12.6) | 0.26 |
| Any animal | 31 | 18 | 4.3 (1.2) | 3.1 (1.6) | 0.01 | 2.8 (1.3) | 2.0 (1.1) | 0.05 | 12.6 (25.8) | 6.2 (15.3) | 0.27 |
| **Animals ever roam free in compound:** | | | | | | | | | | | |
| Chicken/duck | 32 | 17 | 4.2 (1.2) | 3.1 (1.7) | 0.02 | 2.7 (1.3) | 2.1 (1.1) | 0.06 | 12.3 (25.4) | 6.4 (15.9) | 0.20 |
| Cattle/buffalo | 14 | 35 | 4.1 (1.3) | 3.7 (1.6) | 0.56 | 2.7 (1.4) | 2.4 (1.2) | 0.58 | 17.3 (35.8) | 7.5 (13.9) | 0.93 |
| Goat/sheep | 21 | 28 | 4.2 (1.4) | 3.5 (1.5) | 0.11 | 2.9 (1.4) | 2.2 (1.1) | 0.10 | 15.3 (30.4) | 6.5 (13.3) | 0.61 |
| Any animal | 36 | 13 | 4.2 (1.3) | 2.9 (1.6) | 0.01 | 2.7 (1.3) | 1.9 (0.9) | 0.04 | 11.3 (4.0) | 7.6 (18.6) | 0.35 |
| **Animals kept inside home at night:** | | | | | | | | | | | |
| Chicken/duck | 2 | 27 | 3.2 (1.1) | 4.4 (1.3) | 0.20 | 2.5 (0.6) | 2.8 (1.4) | 0.80 | 50.7 (69.8) | 10.6 (21.2) | 0.28 |
| Cattle/buffalo | 3 | 21 | 5.6 (0.5) | 4.0 (1.4) | 0.05 | 4.2 (1.2) | 2.7 (1.2) | 0.09 | 11.0 (15.1) | 17.8 (31.9) | 0.40 |
| Goat/sheep | 8 | 13 | 5.1 (0.8) | 3.6 (2.4) | 0.02 | 3.5 (1.3) | 2.5 (1.4) | 0.19 | 5.9 (9.9) | 20.4 (37.4) | 0.72 |
| Any animal | 9 | 26 | 4.8 (1.2) | 3.9 (1.4) | 0.08 | 3.3 (1.2) | 2.6 (1.3) | 0.15 | 16.4 (32.7) | 12.4 (23.4) | 0.48 |
| **Animal feces observed on floor:** | | | | | | | | | | | |
| Chicken/duck | 2 | 47 | 5.9 (0.0) | 3.7 (1.5) | 0.03 | 5.2 (0.1) | 2.4 (1.2) | 0.02 | 23.7 (6.6) | 9.9 (23.2) | 0.06 |
| Cattle/buffalo | 2 | 47 | 4.5 (0.7) | 3.8 (1.5) | 0.65 | 2.5 (0.6) | 2.5 (1.3) | 0.88 | 1.1 (0.2) | 10.9 (23.3) | 0.96 |
| Goat/sheep | 1 | 48 | 5.9 (--) | 3.8 (1.5) | 0.13 | 5.3 (--) | 2.4 (1.2) | 0.09 | 28.4 (--) | 10.1 (23.0) | 0.16 |
| Any animal | 4 | 45 | 5.4 (0.9) | 3.7 (1.5) | 0.02 | 4.0 (1.3) | 2.4 (1.2) | 0.03 | 6.6 (8.3) | 10.9 (23.9) | 0.25 |

SD: Standard deviation; MW: Mann-Whitney U-test

**Table S5.** Number of animals owned by quartiles of generic and cefotaxime-resistant *E. coli* counts and percent abundance of cefotaxime-resistant *E. coli* on floor swabs

| **Abundance quartile** | | **Chicken/duck** | **Cattle/buffalo** | **Goat/sheep** | **Any animal** |
|---|---|---|---|---|---|
| **Owned by household** | **N** | **Mean (SD)** | **Mean (SD)** | **Mean (SD)** | **Mean (SD)** |
| Generic *E. coli* | | | | | |
| Non-detect | 3 | 0.0 (0.0) | 0.0 (0.0) | 0.0 (0.0) | 0.0 (0.0) |
| Bottom tertile (1.5-3.3) | 16 | 9.1 (16.1) | 1.5 (2.7) | 2.1 (4.5) | 12.7 (20.6) |
| Middle tertile (3.5-4.8) | 15 | 10.4 (13.6) | 1.1 (1.9) | 1.1 (2.0) | 12.6 (15.6) |
| Top tertile (4.9-5.9) | 15 | 21.8 (37.1) | 2.8 (2.5) | 2.3 (2.2) | 26.9 (39.1) |
| Trend test p-value | | 0.01 | 0.02 | 0.05 | 0.004 |
| Cefotaxime-resistant *E. coli* | | | | | |
| Non-detect | 15 | 5.5 (8.5) | 1.2 (2.3) | 1.1 (2.1) | 7.9 (11.8) |
| Bottom tertile (1.5-2.7) | 12 | 9.5 (17.5) | 0.6 (1.2) | 2.2 (4.9) | 12.3 (21.9) |
| Middle tertile (2.7-3.4) | 11 | 14.5 (12.9) | 2.7 (3.1) | 1.3 (2.1) | 18.5 (13.9) |
| Top tertile (3.5-5.3) | 11 | 24.7 (43.8) | 2.4 (2.4) | 2.5 (2.5) | 29.6 (49.6) |
| Trend test p-value | | 0.02 | 0.04 | 0.10 | 0.01 |
| Percent abundance | | | | | |
| Zero | 12 | 6.9 (9.0) | 1.6 (2.5) | 1.4 (2.3) | 9.9 (12.5) |
| Bottom tertile (0.1-1.3%) | 12 | 9.5 (9.6) | 2.0 (2.6) | 1.3 (1.4) | 12.8 (11.7) |
| Middle tertile (1.5-6.3%) | 11 | 17.7 (20.1) | 1.4 (2.2) | 2.8 (5.3) | 21.9 (24.6) |
| Top tertile (8.9-100%) | 11 | 21.5 (44.3) | 2.2 (2.7) | 2.0 (2.5) | 25.6 (50.0) |
| Trend test p-value | | 0.33 | 0.48 | 0.52 | 0.22 |
| **Owned by compound** | | **Mean (SD)** | **Mean (SD)** | **Mean (SD)** | **Mean (SD)** |
| Generic *E. coli* | | | | | |
| Non-detect | 3 | 0.0 (0.0) | 0.0 (0.0) | 2.7 (4.6) | 2.7 (4.6) |
| Bottom tertile (1.5-3.3) | 16 | 9.8 (16.1) | 2.0 (3.7) | 2.1 (4.5) | 14.0 (21.2) |
| Middle tertile (3.5-4.8) | 15 | 12.9 (13.4) | 2.2 (2.4) | 1.3 (2.1) | 16.4 (14.8) |
| Top tertile (4.9-5.9) | 15 | 22.7 (37.2) | 2.9 (2.6) | 2.1 (2.0) | 27.7 (39.1) |
| Trend test p-value | | 0.01 | 0.02 | 0.25 | 0.01 |
| Cefotaxime-resistant *E. coli* | | | | | |
| Non-detect | 15 | 5.6 (9.2) | 1.9 (3.6) | 1.7 (2.7) | 9.1 (13.7) |
| Bottom tertile (1.5-2.7) | 12 | 12.5 (17.2) | 1.5 (1.8) | 2.2 (4.9) | 16.2 (21.1) |
| Middle tertile (2.7-3.4) | 11 | 15.7 (14.1) | 2.7 (3.1) | 1.3 (2.1) | 19.7 (15.5) |
| Top tertile (3.5-5.3) | 11 | 25.8 (43.3) | 3.0 (2.8) | 2.5 (2.1) | 31.4 (45.3) |
| Trend test p-value | | 0.02 | 0.08 | 0.14 | 0.01 |
| Percent abundance | | | | | |
| Zero | 12 | 7.0 (9.8) | 2.3 (3.9) | 1.4 (2.3) | 10.8 (14.9) |
| Bottom tertile (0.1-1.3%) | 12 | 12.8 (11.8) | 2.3 (2.5) | 1.3 (1.4) | 16.3 (13.7) |
| Middle tertile (1.5-6.3%) | 11 | 18.7 (19.3) | 2.1 (2.4) | 2.8 (5.3) | 23.6 (23.2) |
| Top tertile (8.9-100%) | 11 | 22.5 (43.9) | 2.8 (3.0) | 2.0 (2.1) | 27.4 (47.9) |
| Trend test p-value | | 0.21 | 0.43 | 0.34 | 0.11 |

**Table S6.** Prevalence and log10-transformed most probable number (MPN) of *E. coli* on child hands by animal ownership and management practices

| | N | | Prevalence % (n) | | Chi2 | Log10 MPN Mean (SD) | | MW |
|---|---|---|---|---|---|---|---|---|
| | Yes | No | Yes | No | p-value | Yes | No | p-value |
| Soil floors | 19 | 17 | 57.9 (11) | 47.1 (8) | 0.52 | 1.6 (1.1) | 1.0 (0.9) | 0.14 |
| **Household owns at least 1:** | | | | | | | | |
| Chicken/duck | 20 | 16 | 60.0 (12) | 43.8 (7) | 0.33 | 1.5 (1.0) | 1.1 (1.1) | 0.24 |
| Cattle/buffalo | 16 | 20 | 68.8 (11) | 40.0 (8) | 0.09 | 1.6 (1.0) | 1.1 (1.0) | 0.08 |
| Goat/sheep | 12 | 24 | 66.7 (8) | 45.8 (11) | 0.24 | 1.5 (1.0) | 1.2 (1.0) | 0.26 |
| Any animal | 24 | 12 | 58.3 (14) | 41.7 (5) | 0.35 | 1.3 (1.0) | 1.2 (1.2) | 0.51 |
| **Compound owns at least 1:** | | | | | | | | |
| Chicken/duck | 22 | 14 | 59.1 (13) | 42.9 (6) | 0.34 | 1.4 (1.0) | 1.1 (1.1) | 0.27 |
| Cattle/buffalo | 20 | 16 | 65.0 (13) | 37.5 (6) | 0.10 | 1.5 (1.0) | 1.0 (1.1) | 0.10 |
| Goat/sheep | 14 | 22 | 57.1 (8) | 50.0 (11) | 0.68 | 1.4 (1.0) | 1.3 (1.1) | 0.67 |
| Any animal | 27 | 9 | 55.6 (15) | 44.4 (4) | 0.56 | 1.3 (0.9) | 1.3 (1.3) | 0.82 |
| **Animals ever roam free in household:** | | | | | | | | |
| Chicken/duck | 21 | 15 | 57.1 (12) | 46.7 (7) | 0.54 | 1.4 (1.0) | 1.1 (1.1) | 0.29 |
| Cattle/buffalo | 7 | 29 | 71.4 (5) | 48.3 (14) | 0.27 | 1.7 (0.9) | 1.2 (1.0) | 0.17 |
| Goat/sheep | 10 | 26 | 70.0 (7) | 46.2 (12) | 0.20 | 1.6 (1.0) | 1.2 (1.0) | 0.16 |
| Any animal | 22 | 14 | 59.1 (13) | 42.9 (6) | 0.34 | 1.4 (1.0) | 1.1 (1.1) | 0.27 |
| **Animals ever roam free in compound:** | | | | | | | | |
| Chicken/duck | 23 | 13 | 56.5 (13) | 46.2 (6) | 0.55 | 1.4 (1.0) | 1.2 (1.1) | 0.43 |
| Cattle/buffalo | 8 | 28 | 75.0 (6) | 46.4 (13) | 0.15 | 1.5 (0.9) | 1.2 (1.1) | 0.30 |
| Goat/sheep | 12 | 24 | 58.3 (7) | 50.0 (12) | 0.64 | 1.4 (1.1) | 1.2 (1.0) | 0.50 |
| Any animal | 25 | 11 | 56.0 (14) | 45.5 (5) | 0.56 | 1.3 (1.0) | 1.3 (1.2) | 0.65 |
| **Animals kept inside home at night [a]:** | | | | | | | | |
| Chicken/duck | 0 | 20 | -- | 60.0 (12) | -- | -- | 1.5 (1.0) | -- |
| Cattle/buffalo | 1 | 15 | 100.0 (1) | 73.3 (11) | -- | 0.4 (--) | 1.7 (1.0) | -- |
| Goat/sheep | 4 | 8 | 25.0 (1) | 87.5 (7) | -- | 0.9 (1.0) | 1.8 (0.3) | -- |
| Any animal | 4 | 20 | 25.0 (1) | 65.0 (13) | -- | 0.9 (1.0) | 1.4 (0.9) | -- |
| **Animal feces observed on floor [a]:** | | | | | | | | |
| Chicken/duck | 1 | 35 | 0.0 (0) | 54.3 (19) | -- | 0.4 (--) | 1.3 (1.0) | -- |
| Cattle/buffalo | 0 | 36 | -- | 52.8 (19) | -- | -- | 1.3 (1.0) | -- |
| Goat/sheep | 0 | 36 | -- | 52.8 (19) | -- | -- | 1.3 (1.0) | -- |
| Any animal | 3 | 33 | 33.3 (1) | 54.6 (18) | -- | 1.1 (1.2) | 1.3 (1.0) | -- |

SD: Standard deviation; MW: Mann-Whitney U-test.

[a] No statistical test performed because of small cells.

**Table S7.** Number of animals owned by quartiles of *E. coli* counts on child hands

| **Abundance quartile** | | **Chicken/duck** | **Cattle/buffalo** | **Goat/sheep** | **Any animal** |
|---|---|---|---|---|---|
| **Owned by household** | **N** | **Mean (SD)** | **Mean (SD)** | **Mean (SD)** | **Mean (SD)** |
| Generic *E. coli* | | | | | |
| Non-detect | 17 | 5.8 (8.7) | 0.8 (1.5) | 0.8 (1.8) | 7.4 (11.4) |
| Bottom tertile (1.0-1.6) | 7 | 18.0 (24.6) | 1.1 (1.9) | 3.3 (6.3) | 22.4 (29.8) |
| Middle tertile (1.9-2.3) | 6 | 14.5 (11.8) | 1.5 (1.9) | 1.7 (2.9) | 17.7 (14.6) |
| Top tertile (2.4-3.8) | 6 | 40.0 (56.2) | 3.8 (3.5) | 2.0 (2.4) | 45.5 (59.3) |
| Trend test p-value | | 0.03 | 0.02 | 0.22 | 0.06 |
| **Owned by compound** | **N** | **Mean (SD)** | **Mean (SD)** | **Mean (SD)** | **Mean (SD)** |
| Generic *E. coli* | | | | | |
| Non-detect | 17 | 5.6 (8.3) | 0.9 (1.5) | 1.5 (2.5) | 8.1 (10.9) |
| Bottom tertile (1.0-1.6) | 7 | 18.0 (24.6) | 1.1 (1.9) | 3.3 (6.3) | 22.4 (29.8) |
| Middle tertile (1.9-2.3) | 6 | 20.5 (6.4) | 3.2 (1.9) | 1.7 (2.9) | 25.3 (6.8) |
| Top tertile (2.4-3.8) | 6 | 42.0 (55.5) | 4.2 (3.6) | 1.5 (1.6) | 47.7 (58.4) |
| Trend test p-value | | 0.01 | 0.01 | 0.65 | 0.02 |

**Table S8.** *E. coli* prevalence and abundance on floor swabs by frequency of animal roaming

| **Animals roaming frequency** | | **Generic *E. coli*** | | **Cefotaxime-resistant *E. coli*** | | |
|---|---|---|---|---|---|---|
| **In household** | **N** | **Prevalence % (n)** | **Log10 MPN Mean (SD)** | **Prevalence % (n)** | **Log10 MPN Mean (SD)** | **% abundance Mean (SD)** |
| Chicken/duck | | | | | | |
| Never | 19 | 84.2 (16) | 3.1 (1.6) | 52.6 (10) | 2.0 (1.1) | 6.2 (14.8) |
| Sometimes | 15 | 100.0 (15) | 4.0 (1.2) | 86.7 (13) | 2.6 (1.1) | 14.9 (27.0) |
| Always | 15 | 100.0 (15) | 4.6 (1.2) | 73.3 (11) | 3.0 (1.5) | 10.7 (26.0) |
| Trend test p-value | | 0.05 | 0.003 | 0.16 | 0.04 | 0.78 |
| Cattle/buffalo | | | | | | |
| Never | 36 | 91.7 (33) | 3.7 (1.6) | 63.9 (23) | 2.4 (1.3) | 6.6 (13.3) |
| Sometimes | 12 | 100.0 (12) | 4.2 (1.2) | 83.3 (10) | 2.7 (1.3) | 21.8 (38.0) |
| Always | 1 | 100.0 (1) | 5.9 (--) | 100.0 (1) | 4.4 (--) | 3.5 (--) |
| Trend test p-value | | 0.31 | 0.14 | 0.16 | 0.20 | 0.33 |
| Goat/sheep | | | | | | |
| Never | 32 | 90.6 (29) | 3.5 (1.5) | 62.5 (20) | 2.2 (1.1) | 6.0 (12.6) |
| Sometimes | 15 | 100.0 (15) | 4.5 (1.2) | 86.7 (13) | 3.1 (1.3) | 20.3 (34.8) |
| Always | 2 | 100.0 (2) | 4.3 (2.2) | 50.0 (1) | 2.8 (2.3) | 1.8 (2.5) |
| Trend test p-value | | 0.23 | 0.05 | 0.32 | 0.07 | 0.43 |
| Any animal | | | | | | |
| Never | 18 | 83.3 (15) | 3.1 (1.6) | 50.0 (9) | 2.0 (1.1) | 6.2 (15.3) |
| Sometimes | 30 | 100.0 (30) | 4.2 (1.2) | 80.0 (24) | 2.7 (1.3) | 12.9 (26.1) |
| Always | 1 | 100.0 (1) | 5.9 (--) | 100.0 (1) | 4.4 (--) | 3.5 (--) |
| Trend test p-value | | 0.03 | 0.007 | 0.02 | 0.03 | 0.23 |
| **In compound** | **N** | **Prevalence % (n)** | **Log10 MPN Mean (SD)** | **Prevalence % (n)** | **Log10 MPN Mean (SD)** | **% abundance Mean (SD)** |
| Chicken/duck | | | | p=0.05 | | |
| Never | 17 | 82.4 (14) | 3.1 (1.7) | 47.1 (8) | 2.1 (1.1) | 6.4 (15.9) |
| Sometimes | 12 | 100.0 (12) | 4.0 (1.2) | 83.3 (10) | 2.7 (1.3) | 16.8 (30.0) |
| Always | 20 | 100.0 (12) | 4.4 (1.3) | 80.0 (16) | 2.7 (1.3) | 9.6 (22.6) |
| Trend test p-value | | 0.03 | 0.01 | 0.04 | 0.10 | 0.32 |
| Cattle/buffalo | | | | p=0.10 | | |
| Never | 35 | 91.4 (32) | 3.7 (1.6) | 71.4 (25) | 2.4 (1.2) | 7.5 (13.9) |
| Sometimes | 9 | 100.0 (9) | 3.6 (1.0) | 44.4 (4) | 2.2 (1.3) | 12.0 (33.0) |
| Always | 5 | 100.0 (5) | 4.9 (1.4) | 100.0 (5) | 3.8 (1.1) | 26.7 (42.6) |
| Trend test p-value | | 0.30 | 0.22 | 0.71 | 0.17 | 0.51 |
| Goat/sheep | | | | p=0.70 | | |
| Never | 28 | 89.3 (25) | 3.5 (1.5) | 64.3 (18) | 2.2 (1.1) | 6.5 (13.3) |
| Sometimes | 11 | 100.0 (11) | 4.0 (1.5) | 72.7 (8) | 2.8 (1.6) | 16.2 (31.6) |
| Always | 10 | 100.0 (10) | 4.4 (1.3) | 80.0 (8) | 3.0 (1.3) | 14.2 (30.6) |
| Trend test p-value | | 0.16 | 0.08 | 0.34 | 0.09 | 0.42 |
| Any animal | | | | p=0.07 | | |
| Never | 13 | 76.9 (10) | 2.9 (1.6) | 46.2 (6) | 1.9 (0.9) | 7.6 (18.6) |
| Sometimes | 32 | 100.0 (32) | 4.1 (1.3) | 75.0 (24) | 2.6 (1.4) | 9.4 (19.8) |
| Always | 4 | 100.0 (4) | 4.6 (1.5) | 100.0 (4) | 3.4 (0.7) | 26.3 (49.2) |
| Trend test p-value | | 0.01 | 0.01 | 0.02 | 0.02 | 0.21 |

**Table S9.** *E. coli* prevalence and abundance on child hands by frequency of animal roaming

| Animals roaming frequency | In household | | | In compound | | |
|---|---|---|---|---|---|---|
| | N | Prevalence % (n) | Log10 MPN Mean (SD) | N | Prevalence % (n) | Log10 MPN Mean (SD) |
| Chicken/duck | | | | | | |
| Never | 15 | 47.7 (7) | 1.1 (1.1) | 13 | 46.2 (6) | 1.2 (1.1) |
| Sometimes | 9 | 55.6 (5) | 1.5 (1.1) | 7 | 57.1 (4) | 1.6 (1.2) |
| Always | 12 | 58.3 (7) | 1.4 (0.9) | 16 | 56.3 (9) | 1.3 (0.9) |
| Trend test p-value | | 0.55 | 0.34 | | 0.60 | 0.59 |
| Cattle/buffalo | | | | | | |
| Never | 29 | 48.3 (14) | 1.2 (1.0) | 28 | 46.4 (13) | 1.2 (1.1) |
| Sometimes | 6 | 83.3 (5) | 1.9 (0.8) | 6 | 83.3 (5) | 1.6 (0.8) |
| Always | 1 | 0.0 (0) | 0.4 (--) | 2 | 50.0 (1) | 1.4 (1.4) |
| Trend test p-value | | 0.59 | 0.42 | | 0.31 | 0.40 |
| Goat/sheep | | | | | | |
| Never | 26 | 46.2 (12) | 1.2 (1.0) | 24 | 50.0 (12) | 1.2 (1.0) |
| Sometimes | 8 | 75.0 (6) | 1.8 (1.1) | 5 | 100.0 (5) | 1.4 (1.4) |
| Always | 2 | 50.0 (1) | 0.9 (0.6) | 7 | 100.0 (7) | 1.5 (0.9) |
| Trend test p-value | | 0.34 | 0.39 | | 0.42 | 0.43 |
| Any animal | | | | | | |
| Never | 14 | 42.9 (6) | 1.1 (1.1) | 11 | 45.5 (5) | 1.3 (1.2) |
| Sometimes | 21 | 61.9 (13) | 1.5 (1.0) | 23 | 56.5 (13) | 1.3 (1.0) |
| Always | 1 | 0.0 (0) | 0.4 (--) | 2 | 50.0 (1) | 1.4 (1.4) |
| Trend test p-value | | 0.60 | 0.48 | | 0.65 | 0.66 |

**Table S10.** *E. coli* prevalence and abundance on floor swabs by animal cohabitation intensity

| | | **Generic *E. coli*** | | **Cefotaxime-resistant *E. coli*** | | |
|---|---|---|---|---|---|---|
| **Animal cohabitation** | **N** | **Prevalence % (n)** | **Log10 MPN Mean (SD)** | **Prevalence % (n)** | **Log10 MPN Mean (SD)** | **% abundance Mean (SD)** |
| Chicken/duck | | | | | | |
| None owned by compound | 18 | 83.3 (15) | 3.1 (1.6) | 44.4 (8) | 2.0 (1.1) | 6.0 (15.4) |
| None owned by household | 20 | 85.0 (17) | 3.2 (1.6) | 55.0 (11) | 2.0 (1.1) | 5.6 (14.4) |
| Kept outside home at night | 27 | 100.0 (27) | 4.4 (1.3) | 77.8 (21) | 2.8 (1.4) | 10.6 (21.2) |
| Kept inside home at night | 2 | 100.0 (2) | 3.2 (1.1) | 100.0 (2) | 2.5 (0.6) | 50.7 (69.8) |
| Trend test p-value | | 0.02 | 0.03 | 0.02 | 0.05 | 0.15 |
| Cattle/buffalo | | | | | | |
| None owned by compound | 20 | 85.0 (17) | 3.4 (1.7) | 55.0 (11) | 2.3 (1.3) | 4.2 (6.0) |
| None owned by household | 25 | 88.0 (22) | 3.5 (1.5) | 60.0 (15) | 2.2 (1.2) | 3.5 (5.5) |
| Kept outside home at night | 21 | 100.0 (21) | 4.0 (1.4) | 76.2 (16) | 2.7 (1.2) | 17.8 (7.0) |
| Kept inside home at night | 3 | 100.0 (3) | 5.6 (0.5) | 100.0 (3) | 4.2 (1.2) | 8.7 (15.1) |
| Trend test p-value | | 0.05 | 0.03 | 0.07 | 0.04 | 0.28 |
| Goat/sheep | | | | | | |
| None owned by compound | 26 | 92.3 (24) | 3.6 (1.5) | 65.4 (17) | 2.2 (1.1) | 6.4 (2.8) |
| None owned by household | 28 | 89.3 (25) | 3.6 (1.5) | 64.3 (18) | 2.2 (1.1) | 6.8 (13.5) |
| Kept outside home at night | 13 | 100.0 (13) | 3.6 (1.4) | 61.5 (8) | 2.5 (1.4) | 20.4 37.4) |
| Kept inside home at night | 8 | 100.0 (8) | 5.1 (0.8) | 100.0 (8) | 3.4 (1.3) | 5.9 (9.9) |
| Trend test p-value | | 0.28 | 0.06 | 0.21 | 0.07 | 0.49 |
| Any animal | | | | | | |
| None owned by compound | 11 | 81.8 (9) | 3.1 (1.6) | 36.4 (4) | 1.8 (0.9) | 0.8 (1.6) |
| None owned by household | 14 | 78.6 (11) | 3.1 (1.6) | 50.0 (7) | 1.8 (0.8) | 1.1 (0.5) |
| Kept outside home at night | 26 | 100.0 (26) | 3.9 (1.4) | 69.2 (18) | 2.6 (1.3) | 12.4 (4.6) |
| Kept inside home at night | 9 | 100.0 (9) | 4.8 (1.2) | 100.0 (9) | 3.3 (1.2) | 16.4 (32.7) |
| Trend test p-value | | 0.02 | 0.01 | 0.01 | 0.01 | 0.04 |

**Table S11.** *E. coli* prevalence and abundance on child hands by animal cohabitation intensity

| | | Generic *E. coli* | |
|---|---|---|---|
| **Animal cohabitation** | **N** | **Prev (n)** | **Mean (SD)** |
| Chicken/duck | | | |
| None owned by compound | 14 | 42.9 (6) | 1.1 (1.1) |
| None owned by household | 16 | 43.8 (7) | 1.1 (1.1) |
| Kept outside home at night | 20 | 60.0 (12) | 1.5 (1.0) |
| Kept inside home at night | 0 | -- | -- |
| Trend test p-value | | 0.42 | 0.31 |
| Cattle/buffalo | | | |
| None owned by compound | 16 | 37.5 (6) | 1.0 (1.1) |
| None owned by household | 20 | 40.0 (8) | 1.1 (1.0) |
| Kept outside home at night | 15 | 73.3 (11) | 1.7 (1.0) |
| Kept inside home at night | 1 | 0.0 (0) | 0.4 (--) |
| Trend test p-value | | 0.14 | 0.13 |
| Goat/sheep | | | |
| None owned by compound | 22 | 50.0 (11) | 1.3 (1.1) |
| None owned by household | 24 | 45.8 (11) | 1.2 (1.0) |
| Kept outside home at night | 8 | 87.5 (7) | 1.8 (0.9) |
| Kept inside home at night | 4 | 25.0 (1) | 0.9 (1.0) |
| Trend test p-value | | 0.73 | 0.69 |
| Any animal | | | |
| None owned by compound | 9 | 44.4 (4) | 1.3 (1.3) |
| None owned by household | 12 | 41.7 (5) | 1.3 (1.0) |
| Kept outside home at night | 20 | 65.0 (13) | 1.4 (0.9) |
| Kept inside home at night | 4 | 25.0 (1) | 0.9 (1.0) |
| Trend test p-value | | 0.88 | 0.97 |

**Table S12.** *E. coli* prevalence and abundance on floor swabs and child hands by cross-categories of household floor type and animal ownership

| | **Soil floor** | | **Concrete floor** | | |
|---|---|---|---|---|---|
| | **Animals** | **No animals** | **Animals** | **No animals** | **p-value** [a] |
| **Floor swabs** | **N = 22** | **N = 6** | **N = 13** | **N = 8** | |
| Generic *E. coli* | | | | | |
| Prevalence, % (n) | 100.0 (22) | 100.0 (6) | 100.0 (13) | 62.5 (5) | 0.001 |
| Log10 MPN, mean (SD) | 4.8 (1.0) | 4.5 (1.0) | 3.1 (1.2) | 2.0 (0.8) | 0.001 |
| Cefotaxime-resistant *E. coli* | | | | | |
| Prevalence, % (n) | 90.9 (20) | 83.3 (5) | 53.9 (7) | 25.0 (2) | 0.003 |
| Log10 MPN, mean (SD) | 3.3 (1.2) | 2.6 (0.8) | 1.9 (1.1) | 1.3 (0.1) | 0.001 |
| Percent abundance [b], mean (SD) | 14.5 (28.8) | 1.3 (1.8) | 11.6 (20.0) | 0.9 (1.4) | 0.19 |
| **Child hands** | **N = 13** | **N = 6** | **N = 11** | **N = 6** | |
| Generic *E. coli* | | | | | |
| Prevalence, % (n) | 53.8 (19) | 66.7 (4) | 63.6 (7) | 16.7 (1) | 0.25 |
| Log10 MPN, mean (SD) | 1.4 (1.0) | 1.9 (1.4) | 1.3 (0.9) | 0.6 (0.4) | 0.24 |

MPN: Most probable number, SD: Standard deviation.

[a] p-value from chi2 test for prevalence and Kruskal-Wallis test for log10 MPN and percent abundance.

[b] Ratio of MPN counts from IDEXX trays with vs. without cefotaxime supplementation for a given sample.
